# Whole Genome Methylation Sequencing via Enzymatic Conversion (EM-seq): Protocol, Data Processing and Analysis

**DOI:** 10.1101/2023.10.07.561361

**Authors:** Nelly N. Olova, Simon Andrews

## Abstract

Whole genome bisulfite sequencing (WGBS) has been the gold standard technique for base resolution analysis of DNA methylation for the last 15 years. It has been, however, associated with technical biases, which lead to overall overestimation of global and regional methylation values, and significant artifacts in extreme cytosine-rich DNA sequence contexts. Enzymatic conversion of cytosine is the newest approach, set to replace entirely the use of the damaging bisulfite conversion of DNA. The EM-seq technique utilises TET2, T4-BGT and APOBEC in a two-step conversion process, where the modified cytosines are first protected by oxidation and glucosylation, followed by deamination of all unmodified cytosines to uracil. As a result, EM-seq is degradation-free and bias-free, requires low DNA input, and produces high library yields with longer reads, little batch variation, less duplication, uniform genomic coverage, accurate methylation over a larger number of captured CpGs, and no sequence-specific artifacts.

## 1. Introduction

DNA methylation is an epigenetic modification with established importance in the study of both healthy development and disease [1]. It is present in various levels and patterns across prokaryotes and eukaryotes including all vertebrates [2], plants [3], some but not all invertebrates including many insects [4], and some fungi [5]. In the last 15 years, the genomic distribution of 5-methyl-cytosine (5mC) has been successfully characterised at single base resolution through the combination of the bisulfite conversion method [6] with high-throughput, next-generation sequencing [7, 8], establishing whole genome bisulfite-sequencing (WGBS) as the gold standard for studying DNA methylation. WGBS has since enabled a comprehensive characterisation of the distribution and dynamics of 5mC in numerous cellular models, tissues, and organisms in fundamental molecular processes and in pathology, which has contributed greatly to our current understanding of its roles.

The original pre-bisulfite library preparation protocol, known as BS-seq or MethylC-seq [9], where DNA is fragmented, end repaired and ligated to a methylated adapter before bisulfite conversion and amplification, required large amounts of DNA (typically 0.5 -4 μg) due to bisulfite-induced DNA degradation [10]. This shortcoming was mitigated with the development of post-bisulfite WGBS protocols [11, 12], where the bisulfite conversion-associated degradation was harnessed to provide the initial fragmentation, and ligation of the adapters after conversion greatly reduced degradation losses. This allowed sequencing of much lower amounts of input DNA, down to single cells [13, 14]. With the ever increasing use of WGBS, however, it became evident that all approaches relying on bisulfite conversion suffer from various shortcomings, such as uneven and biased genomic coverage, skewed base composition and high inter-sample and inter-protocol variability, which resulted in errors in methylation signal output and interpretation [12, 15–19]. More recently, a novel enzymatic methyl-sequencing (EM-seq) approach for the detection of 5mC has been developed [20] to replace the use of the highly destructive chemical conversion through sodium bisulfite, which was recognized as the root cause for the biases and artefacts in WGBS libraries [16]. EM-seq utilizes a combination of enzymatic reactions to convert cytosine to uracil (**Fig. 1**) with similar efficiency to bisulfite conversion, while preserving the DNA backbone and remaining degradation and bias-free [18, 20, 21]. As a result, EM-seq delivers higher library yields with longer reads and highly uniform coverage over all sequence contexts, including the extremely C-rich asymmetric unmethylated regions, which constitute the main source of false positive signals in classical BS-seq libraries due to a combination of high degradation coupled with conversion-resistance [16]. Thus, EM-seq achieves the highest accuracy in methylation estimation, together with very high reproducibility between samples, little batch variation, less duplication and a larger number of captured CpGs [18–22]. These features make it superior to WGBS and suitable for all applications, including low input and fragmented material, such as FFPE, cell-free DNA, and single cells [23]. In particular, EM-seq can resolve cases of elusive and questionably low methylation levels, strongly affected by false positive artefacts from the bisulfite, such as in mitochondrial DNA and fruit fly DNA, which have been the subject of disputes for several decades [24–28].

**Fig. 1:**
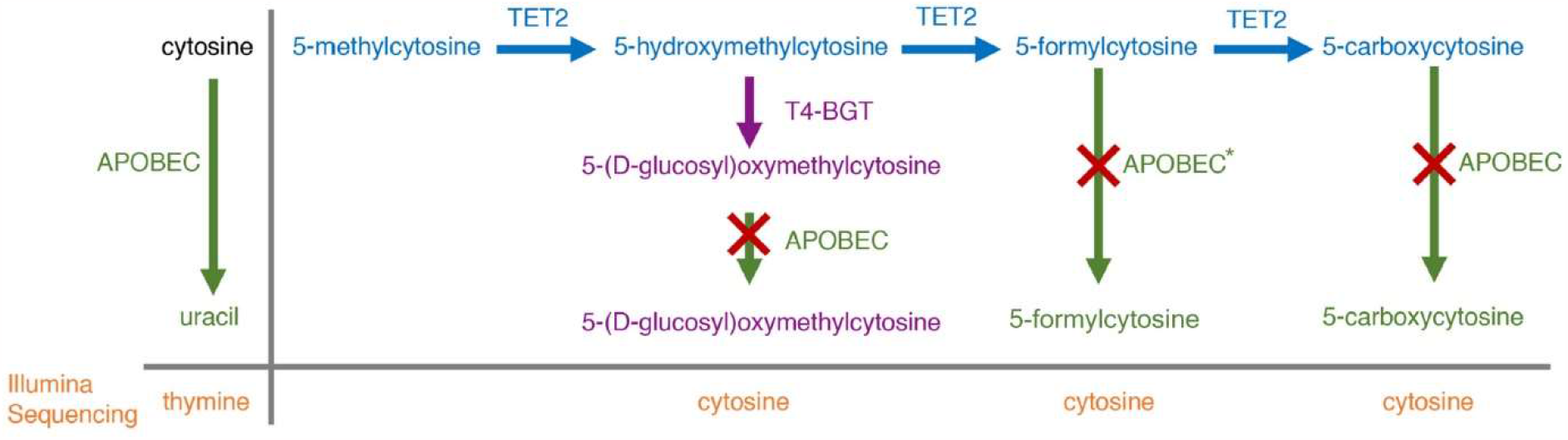
A schematic of the enzymatic conversion of cytosine and its derivatives, reprinted and modified from [20] under CC BY-NC 4.0 licence. First, 5mC is oxidised by TET2 (ten-eleven translocation methylcytosine dioxygenase 2) to 5hmC. 5hmC can be further modified using two possible routes. T4-BGT (T4-phage beta-glucosyltransferase) can glucosylate 5hmC to 5gmC or TET2 can further oxidise 5hmC into 5fC and then 5caC. TET2 and T4-BGT work together so that both enzymes complete the modification of native 5mC and 5hmC to nearly 100%. In a subsequent reaction, APOBEC (apolipoprotein B mRNA editing enzyme catalytic subunit 3A) deaminates unmodified cytosines and converts them into uracils, while 5mC, which has been modified into 5gmC (the major derivative product), 5fC and 5caC, remains resistant to deamination (there might be negligible amount of 5fC with low conversion to 5fU) [20]. *APOBEC deaminates trace amounts of 5fC, and 0 % of 5caC and 5gmC [20].

The current protocol features the New England Biolabs (NEB) whole genome library preparation kit, originally used successfully for the classical BS-seq protocol, and more recently, optimised for enzymatic conversion. The principle and steps are identical to the original BS-seq pre-bisulfite protocol, but enzymatic conversion replaces the bisulfite conversion step (**Fig. 2**, [29]). Therefore, if it is strictly necessary to use bisulfite conversion, such as adding samples to existing public or unpublished BS-seq data, the current protocol is applicable; however, both the bisulfite conversion kit and associated methylated adapters must be purchased separately (*see* **Note 1**). It is important to note that neither WGBS nor EM-seq can distinguish between 5mC and 5-hydroxy-methylcytosine (5hmC) and both modifications appear as positive signals. In EM-seq, unlike WGBS, native 5-formyl-cytosine (5fC) and 5-carboxy-cytosine (5caC) are not converted to uracil and are also read as positive signals (**Fig. 1**, [20]); however, their contribution to the total positive calls pool is negligible.

**Fig. 2:**
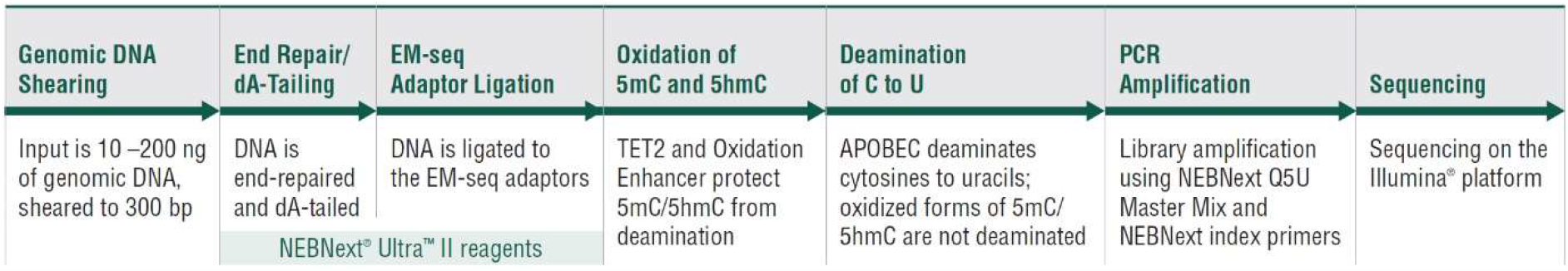
A schematic of the EM-seq workflow. Library preparation steps are performed with the NEBNext Ultra II kit reagents, followed by a two-step enzymatic conversion and library amplification with a uracil-tolerant DNA polymerase (Q5U). Reprinted from www.neb.com (2023) with permission from New England Biolabs, Inc.

An advantage of EM-seq is that the output is identical to WGBS in that unmethylated cytosines appear as thymines, making the analysis pipelines developed for WGBS entirely suitable and directly transferrable to EM-seq. The Bioinformatics Group at the Babraham Institute, Cambridge, were among the first to develop tools for the analysis of WGBS data and in the last 10 years those tools have become some of the best established and highly used worldwide. Each tool has a detailed User Guide and a wealth of associated information on the respective website or GitHub pages. Here we describe the typical DNA methylation processing and analysis workflow and its capabilities for an end-to-end analysis of classical WGBS data, now fully transferrable for data generated by EM-seq.

## 2. Materials

Reagents in this list, where indicated, must be purchased from the recommended suppliers to guarantee the quality of outcome. For equipment and plasticware, the manufacturers are suggestions, and any preferred supplier of good quality consumables or equipment can be used. Please note, it is important to use low retention (i.e., non-stick) nuclease-free plasticware throughout, especially pipette tips.

### 2.1 Reagents and kits

1. Genomic DNA isolated in a manual or automated fashion, excluding chloroform-containing protocols [30]
2. Qubit DNA Quantification Assay kit (High Sensitivity) or Quant-iT PicoGreen dsDNA Assay Kit with Lambda DNA standard (Invitrogen/Thermo Fisher Scientific)
3. NEBNext UltraShear for 24 or 96 reactions (recommended instead of ultra-sonication)
4. NEBNext Enzymatic Methyl-seq Kit for 24 or 96 samples
5. Formamide – recommended, or optional 0.1 N NaOH
6. 80% Ethanol, freshly prepared from 100% ethanol and ultrapure nuclease-free water
7. Low TE buffer (10 mM Tris pH 8.0, 0.1 mM EDTA), or 10 mM Tris pH 8.0 buffer
8. High purity nuclease-free water

### 2.2 Specialist laboratory equipment

1. Qubit fluorometer or a fluorescent microplate reader, such as a PHERAstar FS instrument
2. Covaris ultrasonicator, Bioruptor or other DNA fragmentation instrument capable of handling 50 μl of sample volume and compatible with PCR plates or strip tubes (optional, if preferred instead of UltraShear enzymatic fragmentation, *see* **Note 2**)
3. Magnetic separation rack or plate, such as NEBNext Magnetic Separation Rack or Invitrogen Thermo Fisher DynaMag 96 Side Magnet Well Plate
4. PCR machine
5. Bioanalyzer or TapeStation or other fragment analyzer, together with associated reagents and consumables (always use the ‘high sensitivity’ DNA assays)
6. Benchtop microcentrifuge for 0.2 ml tubes and plates.
7. Vortexer
8. Lab fume hood

### 2.3 Pipettes and plasticware

1. A set of standard pipettes with low retention nuclease-free tips of various sizes
2. Multichannel pipettes: manual 2-20 μl and 20-200 μl 8-channel pipettes (*see* **Note 3**) and an electronic dispensing 100-1200 μl 8-channel pipette with a buffer reservoir
3. Electronic multi-dispenser pipette, such as Eppendorf Multipette E3 with charging adapter and associated combitips of different sizes (*see* **Note 4**)
4. Pipet-aid/pipettor with stripettes
5. 0.2 ml PCR strip tubes (clear) or clear 96-well PCR plates
6. Adhesive seals for PCR plates and seal applicators
7. Plastic tubes of different sizes – 0.5 ml to 5 ml tubes, and 15- or 50-ml conical centrifuge tubes
8. Ice bucket with ice.

### 2.4 Analysis tools and equipment

All listed tools should be installed, configured and ready to use.

1. One linux/unix machine (multi-core recommended) with at least 32GB of RAM
2. FastQC for data quality check: http://www.bioinformatics.babraham.ac.uk/projects/fastqc/
3. FastQ Screen for data purity (contamination) check: https://www.bioinformatics.babraham.ac.uk/projects/fastq_screen/
4. TrimGalore! and CutAdapt [31] for read trimming and Illumina adapter removal: http://www.bioinformatics.babraham.ac.uk/projects/trim_galore/; https://github.com/marcelm/cutadapt
5. Bismark for bisulfite converted read mapping and methylation extraction [32]: http://www.bioinformatics.babraham.ac.uk/projects/bismark/
6. bowtie2 is necessary for bismark read mapping [33]: http://bowtie-bio.sourceforge.net/bowtie2/index.shtml
7. samtools is necessary for bismark to create bam files [34]: http://samtools.sourceforge.net/
8. MultiQC – for a final processing summary report [35]: https://multiqc.info/
9. Nextflow configured with an appropriate executor – for running pipelines [36] (*see* **Note 5**): https://www.nextflow.io/
10. SeqMonk for processed data analysis and visualisation: https://www.bioinformatics.babraham.ac.uk/projects/seqmonk/
11. The above tools need Perl, Python, and Java installed, and R for some SeqMonk functionalities.

## 3. Methods

The NEBNext EM-seq Kit Instruction Manual is very detailed [37] and will be updated through the years. The current protocol includes all steps, highlighting additional important points from the practice and sharing further insights in the **Notes** section. A notable difference is that the current protocol focuses on high-throughput library preparation, for projects with 24 to 96 samples, which are performed manually and not with a liquid handling automated system (i.e., a robot). If less experienced, it is advisable to do 24 to 48 samples at a time, to not risk over-drying the beads during clean up steps.

### 3.1 DNA preparation

1. DNA concentrations must be measured by Qubit or Picogreen dsDNA assay before library preparation, according to respective kit’s manufacturer’s instructions (*see* **Note 6**). Higher concentrations of DNA will need to be diluted first to less than 200 ng/μl.
2. Dilute the controls according to EM-seq user manual. Unmethylated Lambda and methylated pUC19 control DNAs must be added to each sample (*see* **Note 7**), standardly starting at a 1:1,000 mass ratio and going much lower, depending on control genome size and required sequencing depth.
3. Calculate all volumes of DNA and low TE/Tris buffer, including the spike-in unmethylated and methylated controls as per the user manual (*see* **Note 8**). The assay is validated for as little as 10 ng DNA (*see* **Note 9**) and up to 200 ng DNA, made in a 26 μl final volume if using UltraShear, or 50 μl if using ultra-sonication (*see* **Note 2**).
4. Prepare each sample by adding the required volumes of low TE/Tris buffer, methylation controls, and sample DNA (*see* **Note 10**). This step should not be performed in haste and should be allowed plenty of time with a large number of samples.

### 3.2. DNA shearing

1. On ice, prepare a mixture of the UltraShear reaction buffer and UltraShear enzyme mix, according to the number of samples. Add some extra volume to aid with dispensing as shown below. Mix well and distribute equal volumes to an 8-well strip for the multichannel pipette (*see* **Note 3**).

2. With a multichannel pipette, add 18 μl of the UltraShear Master Mix to each sample, mixing thoroughly by pipetting up and down.
3. Place in a preheated Thermocycler with a heated lid (75°C) and choose a program according to the desired fragment size as shown in the UltraShear manual. For a 2 x 150 bp paired end (PE) sequencing, which is becoming the standard, use the following: 15 minutes at 37°C, 15 minutes at 65°C and hold at 4°C (*see* **Note 11**). Continue directly with Step 3.3 End prep.
4. For Covaris/Bioruptor: follow instrument handling instructions for shearing the ready samples – choose 300-400 bp if aiming for a 2 x 150 bp PE sequencing. Carefully transfer the sheared DNA to new nuclease-free PCR tube strips or a plate for the next step (*see* **Notes 12** and **13**).

| REAGENT | 1X RXN VOLUME | 24X RXN VOLUME (27x) | 96X RXN VOLUME (100x) |
| --- | --- | --- | --- |
| UltraShear Reaction Buffer | 14.0 µL | 378 µL | 1400 µL |
| UltraShear | 4.0 µL | 108 µL | 400 µL |

### 3.3 End prep (includes end repair and A-tailing)

1. On ice, prepare a master mix of the end prep reaction according to the number of samples, adding some extra volume to aid with dispensing as shown below. Note that the master mix composition will differ depending on the fragmentation method used in Step 3.2. Mix well and distribute equal volumes to an 8-well strip for the multichannel pipette (*see* **Note 3**).

2. With a multichannel pipette, add 5 μl of the End Prep Master Mix to each sample if UltraShear was used in 3.2, or 10 μl of the End Prep Master Mix if non-enzymatic shearing was used, mixing thoroughly by pipetting up and down.
3. Place in a Thermocycler with a heated lid and run the following program: 30 minutes at 20°C, 30 minutes at 65°C and hold at 4°C. Continue with next step.

|  | UltraShear / Sonication | UltraShear / Sonication | UltraShear / Sonication |
| --- | --- | --- | --- |
| REAGENT | 1X RXN VOLUME | 24X RXN VOLUME (27x) | 96X RXN VOLUME (100x) |
| Ultra II End Prep Reaction Buffer | None / 7.0 µL | None / 189 µL | None / 700 µL |
| 500 mM DTT (from UltraShear kit) | 2.0 µL / None | 54 µL / None | 200 µL / None |
| Ultra II End Prep Enzyme Mix | 3.0 µL / 3.0 µL | 81 µL / 81 µL | 300 µL / 300 µL |

### 3.4 EM-seq adapter ligation

1. On ice, prepare a mixture of the Ligation master mix and Enhancer according to the number of samples. Add some extra volume to aid with dispensing as shown below.

2. Add 2.5 μl of EM-seq Adaptor to each sample.
3. Add 31 μl of the prepared Ligation Master Mix + Enhancer to each sample and mix by pipetting very well to ensure efficient ligation (the mixture is viscous).
4. Incubate at 20°C for 15 minutes in a PCR thermal cycler with the heated lid off, then hold at 4ºC. At this point, the samples can be stored at -20°C overnight or continue with clean-up.

| REAGENT | 1X RXN VOLUME | 24X RXN VOLUME (27x) | 96X RXN VOLUME (100x) |
| --- | --- | --- | --- |
| Ligation Enhancer | 1.0 µL | 27 µL | 100 µL |
| Ultra II Ligation Master Mix | 30.0 µL | 810 µL | 3000 µL (3.0 mL) |

### 3.5 Clean-up of adapter ligated DNA

1. Remove Sample Purification Beads from storage and vortex until homogenized.
2. Dispense 110 μl beads to each sample (∼1.18X sonication, ∼1.3X UltraShear), mix by pipetting very well with care, making sure no beads are lost (the wells are full).
3. Incubate the samples with the beads for at least 5 minutes at RT.
4. Prepare a fresh stock of 80% ethanol, using nuclease-free water. Mix by inversion and place at RT. Suggested: 40 ml Et-OH + 10 ml H_2_O (for 96 samples).
5. Transfer sample strip tubes/plate to a magnetic separation rack/plate and incubate at RT until the solution is completely clear of beads.
6. Keeping the tubes/plate on the magnet, carefully remove the supernatant and discard without disturbing the beads, using a manual multichannel pipette set at 190 μl.
7. With the tubes/plate still on the magnet, add 200 μl of 80% ethanol to the wells without disturbing the bead pellet. This step is performed with a dispensing multi-channel pipette going up to 1200 μl (i.e., for 48 samples or half a plate at a time).
8. Quickly remove and discard the ethanol while keeping the samples on the magnet, avoiding aspiration of the bead pellet (with a manual multichannel pipette set at 200 μl). This step should be performed within 2 minutes (i.e., 10 seconds per PCR strip/plate row).
9. Repeat Steps 7 and 8, to perform two 80% ethanol washes in total. Remove as much of the final wash as possible, once with 200 μl tips and a second time with 10 μl tips to remove all drops around the well walls and bottom, without disturbing the bead pellet.
10. Air dry the bead pellets for 2 minutes at RT. The wells completed first in Step 9 will dry first and may be ready for Step 11 by the time the last wells in Step 9 are completed. The beads should be matte or still glossy, with no ethanol drops visible in the tube.
11. Remove the tubes/plate from the magnet and quickly add 29 μl of Elution Buffer to the bead pellet. A multi-dispenser is most suitable to use, to avoid wells with visible ethanol while not letting others over-dry (*see* **Note 4**). Mix thoroughly with a manual multichannel pipette ensuring all beads are off the walls and well resuspended. It is critical to *not* substitute the provided Elution Buffer with TE or another EDTA-containing buffer (*see* **Note 14**).
12. Incubate at RT for at least 1 minute to let DNA elute from the beads. Centrifuge briefly to collect solution at the bottom of the tubes.
13. Transfer the tubes/plate back to the magnet and incubate at RT for at least 5 minutes or until the solution of beads is completely clear.
14. Carefully remove 28 μl of the eluate (with a manual multichannel), ensuring absolutely no beads are carried over and transfer to a new microcentrifuge plate or PCR tubes (*see* **Note 15**). Samples can now be stored at -20°C if needed.

### 3.6 Oxidation of 5mC and 5hmC

1. Prepare TET2 Buffer, depending on the kit purchased. For 24 samples, add 100 μl of TET2 Reaction Buffer to each of the three provided tubes of TET2 Reaction Buffer Supplement (each one is enough for 10 samples), and 400 μl Buffer to each provided Supplement tube for 96 samples (each one is enough for 40 samples). Vortex and spin down, store at -20°C and use within 4 months.
2. Prepare the oxidation reaction master mix by combining all components as shown below and vortex quickly.

3. Add 17 μl of the Oxidation Master Mix to each sample and mix well by pipetting.
4. Freshly prepare the Fe(II) solution by adding 1 μl of 500 mM Fe(II) stock solution to 1249 μl of water.
5. Add 5 μl of the diluted 0.4 mM Fe(II) solution to each sample and mix by pipetting very well (*see* **Note 16**). Discard the remaining 0.4 mM Fe(II) solution after use.
6. Incubate at 37ºC for 1 hour in a thermal cycler with a heated lid on (≥ 45°C), then hold at 4ºC.
7. Transfer the samples on ice and add 1 μl of Stop Reagent to each reaction.
8. Incubate again at 37ºC for 30 minutes, in a thermal cycler with a heated lid on (≥ 45°C), then hold at 4ºC. The samples can remain overnight at 4ºC or be transferred to -20ºC.

| REAGENT | 1X RXN VOLUME | 24X RXN VOLUME (27x) | 96X RXN VOLUME (100x) |
| --- | --- | --- | --- |
| TET2 Reaction Buffer + TET2 Reaction Buffer Supplement | 10 μL | 270 μL | 1000 μL (1.0 mL) |
| Oxidation Supplement | 1 μL | 27 μL | 100 μL |
| DTT | 1 μL | 27 μL | 100 μL |
| Oxidation Enhancer | 1 μL | 27 μL | 100 μL |
| TET2 | 4 μL | 108 μL | 400 μL |

### 3.7 Clean-up of TET2-oxidised DNA

1. Vortex the Sample Purification Beads until homogenized and dispense 90 μl (∼1.8X) to each sample. Mix well by pipetting, making sure no beads are lost.
2. Repeat the rest of the steps as described in 3.5, resuspending the beads in 17 μl of Elution Buffer and eluting 16 μl of sample to transfer to new PCR tubes/plate. Again, it is critical to not carry over any beads with the elution buffer (*see* **Note 15**).

### 3.8 Denaturation of DNA with formamide (*see* **Note 17**)

1. Pre-heat a PCR thermal cycler to 85°C with the heated lid on (105°C).
2. In a fume hood, add 4 μl formamide to each eluted 16 μl of TET2 oxidized DNA, mix well by pipetting, and centrifuge briefly.
3. Incubate the samples at 85°C for 10 minutes in the pre-heated thermal cycler.
4. Immediately place on ice for 2-3 minutes to ensure DNA has cooled down completely (i.e., is single stranded) before proceeding to the APOBEC conversion step.

### 3.9 APOBEC deamination (conversion of unmodified C to U)

1. On ice, mix the components as shown below to prepare the APOBEC master mix.

2. Add 80 μl of deamination master mix to each of the 20 μl denatured DNA samples, mix well by pipetting and centrifuge to collect all liquid.
3. Incubate at 37ºC for 3 hours and 4ºC afterwards, in a thermal cycler with a heated lid on (≥ 45°C). The samples can remain overnight at 4ºC or be transferred to -20ºC for storage.

| REAGENT | 1X RXN VOLUME | 24X RXN VOLUME (27x) | 96X RXN VOLUME (100x) |
| --- | --- | --- | --- |
| Nuclease-free water | 68 µL | 1836 µL | 6.8 mL (6800 µL) |
| APOBEC Reaction Buffer | 10 µL | 270 µL | 1 mL (1000 µL) |
| BSA | 1 µL | 27 µL | 100 µL |
| APOBEC | 1 µL | 27 µL | 100 µL |

### 3.10 Clean-up of deaminated DNA

The beads become stickier and more difficult to resuspend in this clean-up. Make sure they are not over-dried at the end; air-drying times may be shorter than in previous clean-ups.

1. Vortex the Sample Purification Beads until homogenized and dispense 65 μl (0.65X) to each sample. Mix very well by pipetting, making sure no beads are lost in the tips.
2. Repeat the rest of the steps as described in 3.5., resuspending the beads in 21 μl of Elution Buffer and eluting 20 μl of sample to new PCR tubes/plate. It is critical not to over-dry the beads before adding the Elution Buffer.

### 3.11. PCR amplification

For index pooling refer to Section 3 from the EM-seq Instruction Manual [37].

1. On ice, add 5 μl of EM-seq Index Primer to each of the purified deaminated DNA samples.
2. Dispense 25 μl of Q5U Master Mix to each sample, mix very well by pipetting and centrifuge briefly.
3. Place the samples in a PCR thermal cycler and run the following program: 98ºC for 30 sec, then cycling at 98°C for 10 seconds, 62°C for 30 seconds and 65°C for 60 seconds (for 3 to 8 cycles depending on starting amount, *see* **Note 18**), and a final extension at 65ºC for 5 minutes, followed by a 4ºC hold. Samples can remain overnight at 4ºC or be stored at -20ºC.

### 3.12 Clean-up of amplified DNA libraries

1. Add 50 μl of nuclease-free water to each 50 μl sample and mix well by pipetting.
2. Vortex the Sample Purification Beads until homogenized and dispense 65 μl (0.65X) to each sample. Mix well by pipetting, making sure no beads are lost in the tips.
3. Repeat the rest of the steps as described in 3.5, resuspending the beads in 21 μl of Elution Buffer or, for long term storage, in 1X TE or low TE. Transfer 20 μl of eluate to new PCR tubes/plate.
4. Proceed to quality control (3.12) or store the samples at -20ºC.

### 3.13 Library Quality Control (QC)

1. Depending on the QC instrument used, aliquot 1 to 3 μl of nuclease-free water in a new strip tube/96-well plate according to the number of samples eluted in Step 3.12 (Bioanalyzer is more sensitive and requires the higher dilution factor).
2. Take 1 μl of library with a multichannel pipette from the elution plate, transfer to the QC plate with water and mix well.
3. Follow the Bioanalyzer or Tapestation instructions to measure. The Bioanalyzer is more accurate but is not high-throughput, while Tapestation can run 96-well plates.
4. The resulting trace (library size histogram, see EM-seq Instruction Manual [37]) must be gated from 200 bp to 1200 bp in order to determine library quantity and molarity. Remember to multiply to the dilution factor for obtaining the final library estimates for informing sequencing facilities.

### 3.14 Raw data processing

The main raw data processing steps of whole methylome sequencing are generally the same, irrespective of the protocol used. For the tools discussed here, the classical library preparation approach (including EM-seq) is the ‘default’ mode, meaning no extra options need to be specified, with the only exception being a possible further trimming of reads (*see* **Note 19**). We describe here running raw data processing on Nextflow, a platform for managing and running pipelines [36], which makes it especially easy for wet lab scientists to perform their own data processing, even with limited coding experience (*see* **Note 20**). Babraham Bioinformatics maintain several Nextflow pipelines on the GitHub repository, ‘nf_bisulfite_WGBS’ being designed for classical BS-seq/EM-seq. The pipeline contains the following components (modules), some of which can also be run separately as single program pipelines:

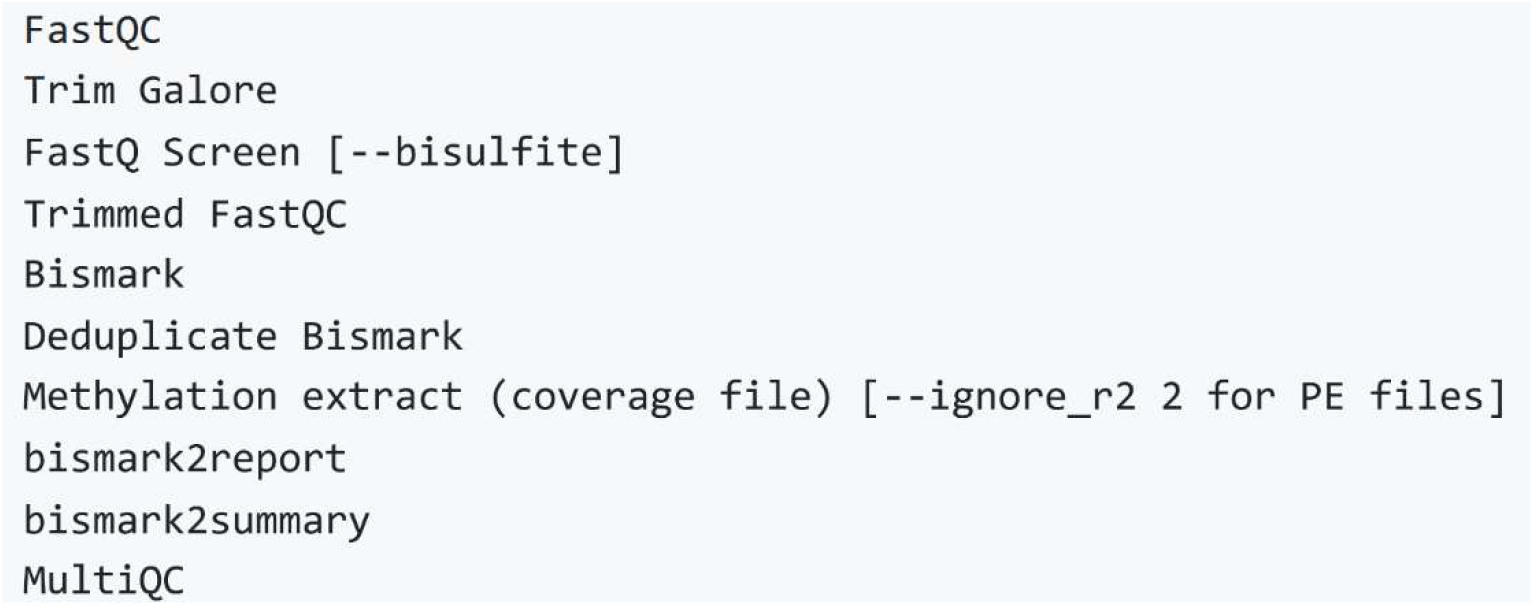

This section includes a breakdown of the analysis steps: the first three are performed by the user, and the rest are performed automatically by the `nf_bisulfite_WGBS` pipeline, unless some changes in steps are required by the user and cannot be added as module arguments as shown in 3.14.4, in which case the separate modules will need to be run independently (not covered here).

#### 3.14.1. Accessing the ‘nf_bisulfite_WGBS’ pipeline

1. Create a fork of Babraham’s Nextflow pipelines repository (https://github.com/s-andrews/nextflow_pipelines) to the user’s own repository.
2. Edit the nextflow.config file to add configurations of the user’s computer system or high-performance cluster (help from local bioinformatics team might be needed). This might also include making sure the individual packages are accessible from the local system.
3. Change the paths to the genomes in the nextflow_pipelines/genomes.d folder with their locations on the user’s system.
4. From a unix command line, run the following on the user’s computer or local cluster (there might be individual systems’ variations):

~~~
module load nextflow
git clone https://github.com/User_profile/nextflow_pipelines.git
cd nextflow_pipelines
git pull
nextflow run nf_bisulfite_WGBS --genome <name-of-genome> *fastq.gz -bg
~~~

where `-bg` stands for running the pipeline non-interactively in the background (see Nextflow @ Babraham User Guide for all details).

#### 3.14.2. Generation of a reference genome

1. The annotated genome of interest must be downloaded in fasta format, where each sequence is annotated with the actual chromosome name and each chromosome sequence is a separate file, such as the searchable Ensembl FTP sequence database: https://www.ensembl.org/info/data/ftp/index.html
2. The most accurate approach is to map simultaneously to the genome of interest and the control DNA sequences via creating a combined genome, rather than aligning separately. To create a combined genome, add pUC19 and Lambda phage sequences as if they were additional chromosomes to the samples’ genome.
3. This step ensures the genomic reference sequence is converted and looks like bisulfite-converted DNA, to make further alignment possible. It is separate from the pipeline because it is performed once per annotated genome. The script takes fasta format sequence files, and after completion, the converted reference genome path must be added to the nextflow_pipelines/genomes.d folder.

~~~
module load bismark
bismark_genome_preparation --path_to_aligner /path/to/bowtie2/ --verbose
/path/to/genome/
~~~

#### 3.14.3. Raw data quality control with FastQC

FastQC provides an overview with summary graphs and tables to help assess data quality, exported in an HTML report and accessible offline. It is best to do this module separately before running the full pipeline, to assess the need for further read trimming (*see* **Note 19**).

nextflow run nf_fastqc *fastq.gz

#### 3.14.4. Trimming of adapters and low-quality bases with TrimGalore!

If FastQC plots show a significant base composition bias at the start and end of your EM-seq library reads (*see* **Note 19**), the pipeline command line should be modified to remove (hard clip) the problematic sequences (the example is for paired end data and 10 bases of biased sequence):

~~~
nextflow run nf_bisulfite_WGBS --genome <name-of-genome> *fastq.gz -bg –
trim_galore_args=“--clip_R1 10 --clip_R2 10 –
three_prime_clip_R1 10 --three_prime_clip_R2 10”
~~~

#### 3.14.5. FastQ Screen and FastQC of trimmed data

1. FastQ Screen scans the data for possible sequence contamination with DNA from another source or species. In `nf_bisulfite_WGBS` it works by default with the argument `--bisulfite` for converted sequences.
2. FastQC performs data quality assessment as above, post-trimming.

#### 3.14.6. Read alignment, deduplication and methylation extraction with Bismark

1. The alignment to a reference genome finds the genomic locations of the read sequences to yield base pair resolution. A very detailed Bismark User Guide (http://felixkrueger.github.io/Bismark/) of how it works is available online.
2. Deduplication is compulsory for WGBS/EM-seq analysis to remove reads with identical coordinates on the same strand, which are considered the result of PCR over-amplification of the same original read.
3. Methylation extraction extracts and compiles all cytosine calls and genomic coordinates from the aligned reads in CG, CHG and CHH contexts (H = A, G or T), and generates lighter files for further analysis. It also removes redundant information from overlapping read 1 (R1) and read 2 (R2) sequences in PE libraries.

#### 3.14.7. Bismark summary reports and MultiQC

1. This step generates per sample reports for the completion of the previous bismark steps, such as aligned and duplicated reads, totals of methylated and unmethylated calls and percentages, and M-bias reports.
2. MultiQC provides summaries for all previously generated reports and logs for all samples and generates an HTML output with plots to visualise overall processing parameters and quality across samples.

### 3.15 Analysis of methylation with SeqMonk

SeqMonk is a program for the analysis of high-throughput sequencing data. It is genome browser-based, with a graphical user interface, making it particularly suitable for wet lab users. There are comprehensive online guides on how SeqMonk works (https://www.bioinformatics.babraham.ac.uk/training.html), and several YouTube tutorials. This chapter focuses on the basics and principles of DNA methylation analysis, broken down in several steps, and further details can be found in the online tutorials and in the SeqMonk ‘Help’ section, which is also comprehensive.

#### 3.15.1. Create a SeqMonk project

You must initially create a project using the same genome assembly version used in step 3.14.2, after which the coverage (.cov) files from the bismark methylation extraction step can be directly imported into the SeqMonk project. Note that in methylation analysis “reads” are the individual CpG calls from the sequencing (read length = 1 bp), and “strand” signifies methylated (“+” and red) or unmethylated (“-” and blue).

#### 3.15.2. Probe generation

In SeqMonk terminology, a “probe” is a region in the genome where we wish to make a quantitative measurement. Probes can be arbitrary tiled regions, or designed around a genomic feature (e.g., genes or CpG islands). Ideally the probes should be designed as windows that cover the entire genome, allowing us to evaluate the data on a global level and facilitating robust comparison between samples. While these windows can be of a fixed base pair length (e.g., 1-5 kb via the ‘Running window probe generator’), because of the varying density of CpGs in the genome, it is better to use windows with a fixed CpG count (i.e., 50-200 CpGs), so that the number of methylation calls in each window is roughly equal. The latter approach is performed with the ‘Read position probe generator’ by selecting all datasets and choosing a preferred CpG count, and results in even coverage with more uniform ‘weights’ of the probes for smoother plots and more robust downstream analyses. The generated probes are stable and will not change until we generate new probes.

#### 3.15.3. Read count quantitation and evaluation

1. First, we count the number of CpG calls identified from our sequencing, on the set of probes we have generated, to evaluate any need for filtering.
2. Check the distribution of reads per probe (read count histogram, **Fig. 3A**), and zoom in if some probes have reads into the 100 000s, as such numbers are clear outliers and most likely mapping artefacts. Since these will affect methylation quantitation, those probes need to be excluded from further analysis.

**Fig. 3:**
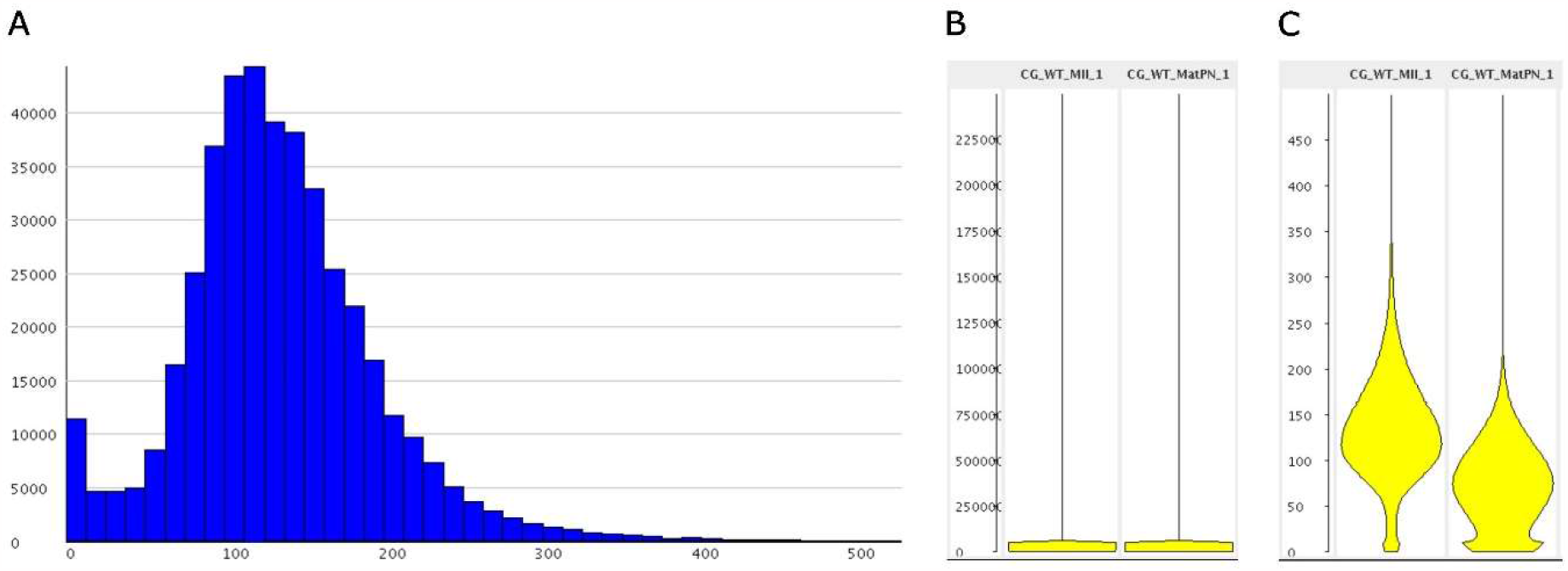
Data filtering prior to methylation analysis based on read count per probe. A) Read count histogram, zoomed in so that high coverage reads are not visible. B) Bean plot of read probe count between samples before filtering and C) after filtering out outliers with a read count above 500.
3. If the probes were generated by fixed number of CpG positions, exploring read length and other metrics compared between the samples, will also give a good idea of the data and where (or for which samples) to expect weaknesses.

#### 3.15.4. Filtering

There are many options for filtering and “filter on value” will allow filtering based on the quantitated value of each probe. Thus, selecting probes with less than 500 read counts as in the given example will create a new probe list where all the high coverage outlier probes are removed (**Fig. 3 B and C**).

#### 3.15.5. Methylation quantitation

The “Bisulfite methylation over features” pipeline (BS-pipeline) is the recommended approach (*see* **Note 21**). It provides data filtering options, such as setting a threshold for minimum CpG observations, i.e., CpG positions covered by at least one call, in order to include a probe (a “horizontal” filtering) and select a minimum depth of coverage for the CpG positions within the probe (a “vertical” filtering). The vertical filtering, although a common practice, has shown a high risk for increasing WGBS coverage biases [16] and should only be used where coverage depth is expected to be extremely high, which is normally associated with targeted methods such as Reduced Representation sequencing (RRBS), and not with whole genome protocols. The horizontal filtering option, however, is highly recommended since probes with fewer than 5 or even 10 CpG positions within the probe are likely to generate unstable methylation percentages and therefore skew results. This data should not be included in further analyses.

#### 3.15.6. Plotting and visualising global methylation levels

Once methylation is quantitated, there are multiple ways to explore and visualise the data.

1. Global 5mC levels are the first thing to assess via a number of ways, such as the bean plot (or violin plot, because it also shows density) or box-whisker and star wars plots, which can visualise global methylation levels across all samples.
2. Histograms show methylation density and distribution, while scatter plots provide a one-to-one comparison between the distributions of two samples, such as a control against a condition or treatment (**Fig. 4A**).

**Fig. 4:**
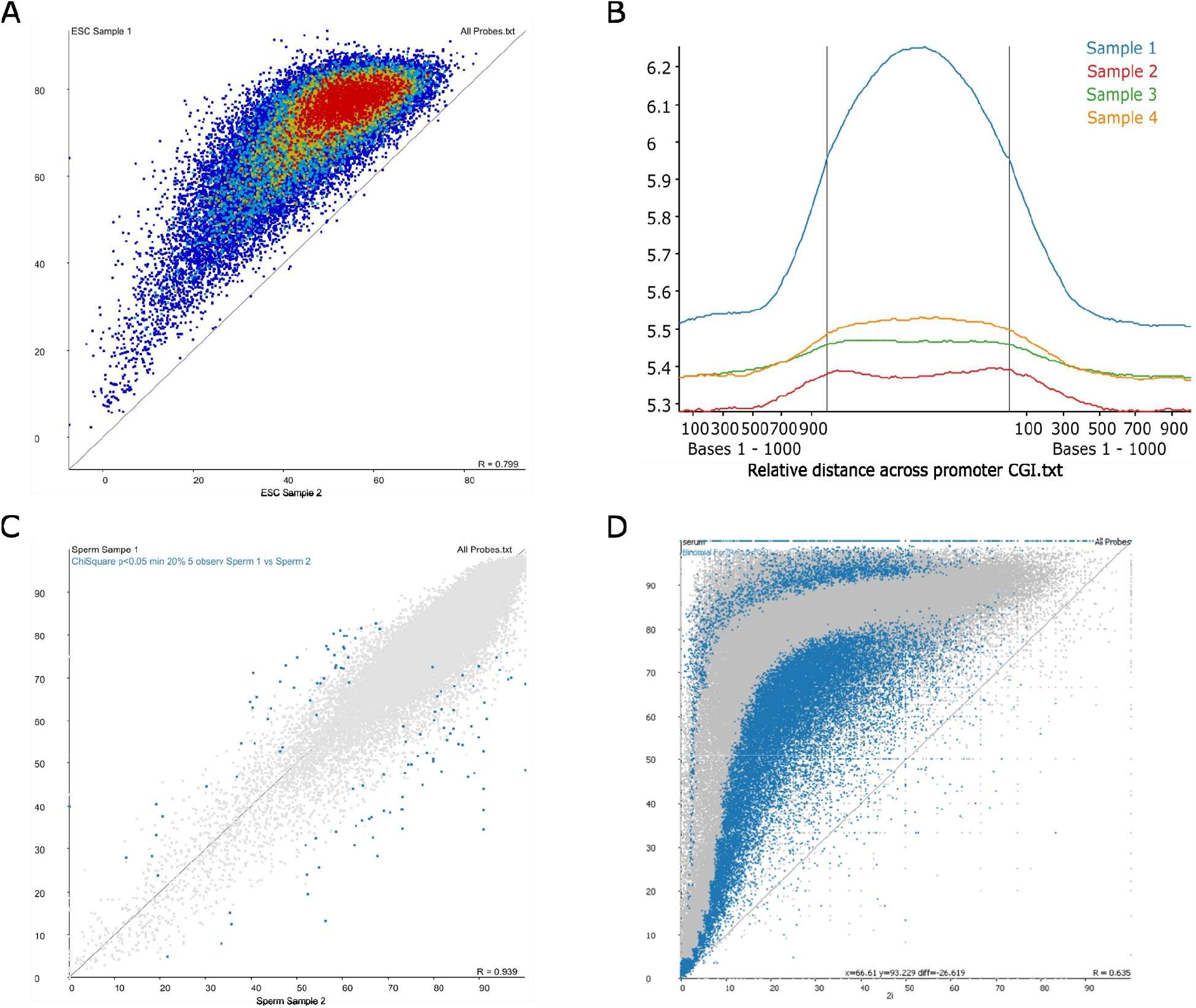
Examples of plots and analyses from SeqMonk. A) Scatter plot, B) A relative trend plot showing methylation levels across a feature (promoter or CGI), C) A chi-square DMR analysis when global methylation levels between samples do not differ, and D) A binomial differential methylation analysis when global methylation levels are significantly different.
3. Other useful options are heat plots, which are a valuable way to look at many samples at once, or the line plots, where methylation levels are shown as a line connecting the different samples. These are useful if looking at time points, stages, or another type of connected process.
4. The plots can be saved in two formats – a PNG for easy viewing and presentations, and a scalable vector SVG format for generating editable publication quality figures.

#### 3.15.7. Analysing genomic features (e.g., genes, promoters, etc.)

1. Feature analysis can be done with the same probes used in the global analysis, by selecting the probes overlapping with a genomic feature (“filter by feature”).
2. The same plots as in 3.15.6 are used for visualisation, but they will only represent methylation of the selected genomic feature, such as promoters, and not the whole genome.
3. A more sophisticated analysis will require the generation of new “feature probes,” which will delete the previously specified probes. Perform quantitation on the new probes as in Step 3.15.5 and visualise as described above.
4. In addition, the averaged methylation trend is commonly investigated over certain genomic features, such as transcriptional start sites (TSS), promoters, CpG islands (CGIs) or gene bodies. Probes are generated to span over the feature, starting before and finishing after for several (kilo)bases. After quantitation, cumulative or relative trend plots are generated (**Fig 4B**).

#### 3.15.8. Assessing differential methylation

This analysis aims to find regions, which are differentially methylated between two samples (i.e., DMRs). It is the most complex, and often most important methylation analysis, involving several statistical considerations. SeqMonk has multiple statistical tests and approaches that can be employed in accordance with the type of data and research question.

1. For data with no global methylation changes between a sample and control, contingency statistics such as chi-square, Fisher’s exact test, logistic regression, or the bisulphite analysis mode of EdgeR [38] are available options for defining thresholds and significance for differential methylation (**Fig 4C**).
2. For datasets where total methylation changes globally from the control, such as during reprogramming stages or other conditions vastly affecting global levels, binomial statistics are more useful in defining regions that change more or less than the global average change. For example, here we look for regions that escape the global changes or such that stand out above the average change (**Fig 4D**).
3. Always check random hits, especially the top hits, for underlying artefacts by zooming in the gene or region location in the SeqMonk browser and visually inspecting if the actual data (methylation calls) make sense.

#### 3.15.9. Generating and saving reports

Reports can be saved at multiple steps of the described analysis workflow.

1. Datastore reports generate summarised information between the different samples, such as total number of methylated and unmethylated cytosines, coverage, and calculated median and mean of global methylation levels.
2. Annotated probe reports contain detailed information about each probe together with the quantitated methylation value and can be imported in R packages for further analysis and visualization.
3. Feature reports are like the annotated reports but summarize data for a specific feature instead.
4. Data can also be exported and saved for the filtered probe lists and from the plots, to be transferred to other packages for visualization or analysis.

## 4. Notes

**Note 1.** The EM-seq protocol is based on the NEBNext Ultra II DNA Library Prep Kit for Illumina, which is offered separately without the Enzymatic Conversion Module and is appropriate for BS-seq. Replace the enzymatic conversion steps described here with bisulfite conversion. It is important to note that the methylated adapters included in the EM-seq kit are different from the methylated adapters used for BS-seq and the latter must be purchased separately from Illumina (see NEBNext Ultra II’s instruction manual). A comprehensive analysis of BS-seq performance with different bisulfite conversion protocols can be found here [16].

**Note 2.** Sonication is a widely used method for DNA shearing, however, there is evidence it preferentially targets methylated DNA and can be used as a hypomethylating agent [39, 40]. NEBNext UltraShear is a novel enzymatic shearing module developed to circumvent the use of ultra-sonication and aid with high-throughput sample fragmentation, being performed in a PCR strip tube or plate format and followed directly by library preparation steps. Enzymatic shearing different from UltraShear is not recommended with the current protocol, as it often leads to loss of DNA methylation (yields short, methylated fragments below retention size).

**Note 3.** Multichannel pipettes are the natural choice for PCR strips and 96-well plates; however, it only works if the tips fit properly on all channels and guarantee pipetting accuracy. It is also important to choose a pipette gentle to the thumbs, as the current high-throughput protocol requires heavy manual pipetting and should not impose a health risk; tip ejection should not require extreme force and cause extra pressure to the thumb (ideally tip ejection should not be performed with the thumb). CAPP and Rainin manual multichannel pipettes are of good quality, CAPP being the multichannel gold standard, also gentle to the hand: its tips fit uniformly and allow very accurate dispensing throughout all channels; tip ejection requires the whole palm. Importantly, the reagent for use with a multichannel pipette needs to be dispensed in an 8-tube strip or a special reservoir in advance, which can be fiddly (200 μl 8-strip tubes may need refills) and usually associated with some master-mix waste. *See* **Note 4** for an alternative.

**Note 4.** Multi-dispensing pipettes (single channel dispensers) can be a very useful alternative to multi-channel pipettes and are much more sparing to the hands, but there is a risk of cross-contamination between samples. This risk can be avoided by aiming the tip at the inside of the well wall, without touching it, so that the drop of liquid sticks to the wall, especially with small volumes. Larger volumes can be ejected directly to the bottom, again without touching anywhere inside the well. Reagent void volume (which is lost inside the dispensing tip) can be very low.

**Notes 5.** The individual workflow components do not need Nextflow and can be run separately. This chapter, however, describes the use of ready-made Nextflow pipelines for WGBS/EM-seq analysis, which is the simplest and most reproducible approach to perform data processing and quality check, and is recommended for publication purposes.

**Note 6.** A Nanodrop spectrophotometer is good for initial quantification but is not appropriate for determining the final DNA concentration for library preparation due to its low accuracy.

**Note 7.** Bisulfite-based analysis traditionally utilized unconverted DNA controls, added to samples in small amounts as a ‘spike-in’ to calculate bisulfite conversion efficiency, while positive controls were not necessary. With the two-step enzymatic conversion, however, an unmethylated control would only provide evidence for the efficacy of the second step, where C is deaminated to U with APOBEC (see **Fig. 1**). A fully methylated control, on the contrary, evidences the efficacy of the TET2 + T4-BGT first reaction, and should appear as fully methylated (i.e., 5mC successfully converted to 5gmC and 5caC) in the post-processing analysis.

**Note 8.** Calculate in MS Excel and print the sample IDs, buffer volume, and DNA sample volume, both as a list and as a 96-well plate/PCR strip schematic, which will greatly reduce pipetting errors. Ordering the samples by increasing or decreasing pipetting volume rather than by sample ID (i.e., buffer decreasing and DNA increasing), will make pipetting easy and quick by not having to make substantial changes to pipet volumes between each sample, and will reduce likelihood of errors, since two neighbouring samples will have the same or similar volumes.

**Note 9.** When using 10 ng DNA with the current protocol, take note that the enzymatic UltraShear fragmentation outperforms ultra-sonication most notably with lower DNA input, achieving significantly higher usable read counts and CpG capture at 10 ng [41]. For even lower starting amounts, EM-seq has been tested and works well with as little as 100 pg. However, some performance parameters will decrease, albeit this has only been tested with ultra-sonication [19, 20]. A specialised low DNA input EM-seq protocol has been optimised by NEB to be released separately.

**Note 10.** Making a pipetting error is easy with many samples, and it is advisable to use a system that will reduce the likelihood of errors. Using the sample list & plate order from **Note 8**, add the low TE/Tris buffer first, then the unmethylated and methylated control DNAs and sample DNA last. Advisably, preparing the conversion controls together with the low TE/Tris buffer as a master mix and dispensing them once altogether, will greatly reduce pipetting effort and possible mistakes, even if it means slightly different sample to control ratios (due to different buffer volume per sample). Double and triple check correct Sample ID and allocated position in plate before pipetting. Cover completed PCR tubes or plate columns to avoid pipetting in the same place.

**Note 11.** EM-seq can handle larger insert sizes and some Illumina instruments can now sequence 600 bp of insert length. The current protocol describes the generation of libraries suitable for 150 bases long PE sequencing (insert size > 300 bp). If smaller size fragments are required, the described protocol applies, with the exception of the final library clean-up where bead to sample ratios would differ: refer to the NEBNext EM-seq or UltraShear Instruction Manuals [37].

**Note 12.** Instructions on ultra-sonication parameters should be provided with the instrument. An example of Covaris S2 sonication to mean fragment size between 350 to 450 bp is: 200 cycles per burst, 10% duty-cycle, intensity of 4 or 5, and treatment time of 50 s (once) or 40 s (twice).

**Note 13.** Low TE/10 mM Tris buffer must be used for sonication and not water, as this helps stabilize and solubilize DNA, mitigating the side effects of biased degradation and loss of methylation from ultra-sonication (*see* **Note 2**).

**Note 14.** EDTA inhibits the TET2 oxidation reaction, therefore the elution at this purification stage is with the provided Elution Buffer, or alternatively with EDTA-free Tris-HCl buffer or nuclease-free water.

**Note 15.** Magnetic beads inhibit the enzymatic reactions that follow and there should be no carryover of beads into the eluate.

**Note 16.** Fe(II) catalyses and starts the TET2 oxidation reaction once all other components are present. Adding Fe(II) earlier with the rest of components may compromise the efficiency of the enzymatic reaction.

**Note 17.** Two options are described in the NEBNext EM-seq Instruction Manual [37]; however, the developers strongly recommend the use of formamide instead of NaOH, therefore only the formamide denaturation is described here. NaOH is more likely to cause problems with concentration if not freshly prepared and yield incompletely denatured DNA, which results in incomplete enzymatic conversion. A detailed protocol for NaOH denaturation can be found in the kit’s instruction manual.

**Note 18.** EM-seq does not need a substantial amplification as classical BS-seq [16, 18], but requires a minimum of 3 PCR cycles to prepare the adapter-ligated DNA fragments with the correct ends for Illumina sequencing [20]. PCR cycle number guide: 3-4 cycles for 100-200 ng DNA input, 5-6 cycles for 50 ng DNA input and 7-8 cycles for 10 ng DNA input.

**Note 19.** There have been observations of base composition bias in EM-seq reads within the first and last 10 bases, both on read 1 (R1) and read 2 (R2), which do not affect cytosine and likely will not affect the methylation call output. However, any remnants of artificially introduced sequence in the reads, which do not belong to the reference genome sequence, could affect downstream mapping. Default alignment parameters are quite stringent, allowing 2 bases mismatch for a 75 base long read, which means such reads can be excluded at the alignment step, leading to a loss of mappable data. While NEB advises removing an extra 5 bases off both R1 and R2, performing quality control with FastQC as a first step will help assess the need for any extra trimming. A conservative approach, especially when working with a large sample size with variable end-bias, would be trimming (i.e., hard clipping) the first and last 10 bases of each read, in addition to the default trimming of low-quality bases and Illumina adapters.

**Note 20.** Nextflow needs to be installed on the user’s local system, such as a high-performance computer cluster, which is an easily executable self-installing procedure (see Nextflow documentation). In addition to the pipeline presented here, Nextflow pipelines are available through the nf-core community framework [42], including ‘methylseq’ (https://nf-co.re/methylseq/2.4.0), which, in its ‘Bismark workflow’ mode, overlaps to a large extent with the workflow described here, and has a parameter `--em_seq` to clip 8 bases of both read ends (R1 & R2) in case of sequence biases. Importantly, nf-core pipelines do not require any prior software installation, they come with pre-built reference genome indices (to download the required genome), and are built with user-friendly web graphical interfaces.

**Note 21.** The Bisulfite methylation pipeline (BS-pipeline) estimates methylation values of each CpG position first by averaging the value from all calls associated with that position, and subsequently averaging the methylation value of all CpGs over the whole probe. The alternative and simplest approach to measure methylation is through “difference quantitation,” where the overall percentage of positive (methylated) calls out of total calls is calculated per probe, irrespective of the CpG position they came from. However, the latter approach was shown to highly bias the methylation estimation in WGBS, since unmethylated CpGs generate overall fewer calls and have lower coverage (i.e., depth per position) than the positive, methylated CpGs [16]. The latter approach will work better with EM-seq; however, this has not been evaluated yet. The BS-pipeline has the added benefit of options for data filtering.

## Acknowledgements

The authors thank Louise Williams, Max Fritsch, and Michael Skelly from New England Biolabs Inc. for valuable discussions, critical reading, and provision of materials, as well as Phil Ewels from Seqera Labs for valuable discussions and critical reading of the manuscript.

## Notes

### Competing Interest Statement

The authors have declared no competing interest.

## References

1. Luo C, Hajkova P, Ecker JR (2018) Dynamic DNA methylation: In the right place at the right time. Science (80-) 361:1336–1340. 10.1126/science.aat6806

2. Schübeler D (2015) Function and information content of DNA methylation. Nature 517:321–326. 10.1038/nature14192

3. Zhang H, Lang Z, Zhu J (2018) Dynamics and function of DNA methylation in plants. Nat Rev Mol Cell Biol 19:489–506. 10.1038/s41580-018-0016-z

4. Regev A, Lamb MJ, Jablonka E (1998) The Role of DNA Methylation in Invertebrates : Developmental Regulation or Genome Defense? Mol Biol Evol 15:880–891

5. Selker EU, Tountas NA, Cross SH, et al (2003) The methylated component of the Neurospora crassa genome. Nature 422:893–897. 10.1038/nature01564

6. Frommer M, Mcdonald LE, Millar DS, et al (1992) A genomic sequencing protocol that yields a positive display of 5-methylcytosine residues in individual DNA strands. PNAS 89:1827–1831

7. Cokus SJ, Feng S, Zhang X, et al (2008) Shotgun bisulphite sequencing of the Arabidopsis genome reveals DNA methylation patterning. Nature 452:215–9. 10.1038/nature06745

8. Lister R, O’Malley RC, Tonti-Filippini J, et al (2008) Highly integrated single-base resolution maps of the epigenome in Arabidopsis. Cell 133:523–36. 10.1016/j.cell.2008.03.029

9. Urich MA, Nery JR, Lister R, et al (2015) MethylC-seq library preparation for base-resolution whole-genome bisulfite sequencing. Nat Protoc 10:475–483. 10.1038/nprot.2014.114

10. Tanaka K, Okamoto A (2007) Degradation of DNA by bisulfite treatment. Bioorg Med Chem Lett 17:1912–1915. 10.1016/j.bmcl.2007.01.040

11. Miura F, Enomoto Y, Dairiki R, Ito T (2012) Amplification-free whole-genome bisulfite sequencing by post-bisulfite adaptor tagging. Nucleic Acids Res 40:e136. 10.1093/nar/gks454

12. Raine A, Manlig E, Wahlberg P, et al (2017) SPlinted Ligation Adapter Tagging (SPLAT), a novel library preparation method for whole genome bisulphite sequencing. Nucleic Acids Res 45:1–15. 10.1093/nar/gkw1110

13. Smallwood SA, Lee HJ, Angermueller C, et al (2014) Single-cell genome-wide bisulfite sequencing for assessing epigenetic heterogeneity. Nat Methods 11:817–820. 10.1038/nmeth.3035

14. Farlik M, Sheffield NC, Nuzzo A, et al (2015) Single-Cell DNA Methylome Sequencing and Bioinformatic Inference of Epigenomic Cell-State Dynamics. Cell Rep 10:1386–1397. 10.1016/j.celrep.2015.02.001

15. Ji L, Sasaki T, Sun X, et al (2014) Methylated DNA is over-represented in wholegenome bisulfite sequencing data. Front Genet 5:1–10. 10.3389/fgene.2014.00341

16. Olova N, Krueger F, Andrews S, et al (2018) Comparison of whole-genome bisulfite sequencing library preparation strategies identifies sources of biases affecting DNA methylation data. Genome Biol 19:1–19. 10.1186/s13059-018-1408-2

17. Zhou L, Ng HK, Drautz-moses DI, et al (2019) Systematic evaluation of library preparation methods and sequencing platforms for highthroughput whole genome bisulfite sequencing. 1–16. 10.1038/s41598-019-46875-5

18. Feng S, Zhong Z, Wang M, Jacobsen SE (2020) Efficient and accurate determination of genome - wide DNA methylation patterns in Arabidopsis thaliana with enzymatic methyl sequencing. Epigenetics Chromatin 1–17. 10.1186/s13072-020-00361-9

19. Foox J, Nordlund J, Lalancette C, et al (2021) The SEQC2 epigenomics quality control (EpiQC) study. Genome Biol 22:1–30. 10.1186/s13059-021-02529-2

20. Vaisvila R, Ponnaluri VKC, Sun Z, et al (2021) Enzymatic methyl sequencing detects DNA methylation at single-base resolution from picograms of DNA. Genome Res 31:1280–1289. 10.1101/gr.266551.120

21. Han Y, Zheleznyakova GY, Marincevic-zuniga Y, et al (2022) Comparison of EM-seq and PBAT methylome library methods for low-input DNA. Epigenetics 17:1195–1204. 10.1080/15592294.2021.1997406

22. Morrison J, Koeman JM, Johnson BK, et al (2021) Evaluation of whole - genome DNA methylation sequencing library preparation protocols. Epigenetics Chromatin 14:1–15. 10.1186/s13072-021-00401-y

23. Chatterton Z, Lamichhane P, Ahmadi Rastegar D, et al (2023) Single-cell DNA methylation sequencing by combinatorial indexing and enzymatic DNA methylation conversion. Cell Biosci 13:1–11. 10.1186/s13578-022-00938-9

24. Nass MMK (1973) Differential Methylation of Mitochondrial and Nuclear DNA in Cultured Mouse, Hamster and Virus-transformed Hamster Cells In vivo and in vitro Methylation. J Mol Biol 155–175

25. van der Wijst MGP, Rots MG (2015) Mitochondrial epigenetics: an overlooked layer of regulation? Trends Genet 31:353–356. 10.1016/j.tig.2015.03.009

26. Shao Z, Han Y, Zhou D (2023) Optimized bisulfite sequencing analysis reveals the lack of 5-methylcytosine in mammalian mitochondrial DNA. BMC Genomics 24:439. 10.1186/s12864-023-09541-9

27. Raddatz G, Guzzardo PM, Olova N, et al (2013) Dnmt2-dependent methylomes lack defined DNA methylation patterns. Proc Natl Acad Sci U S A 110:8627–8631. 10.1073/pnas.1306723110

28. Krauss V, Reuter G (2011) DNA Methylation in Drosophila — A Critical Evaluation, 1st ed. Elsevier Inc.

29. Williams L, Bei Y, Church HE, et al (2019) Enzymatic Methyl-seq: The Next Generation of Methylome Analysis

30. Soriano-Tarraga C, Jimenez-Conde J, Giralt-Steinhauer E, et al (2013) DNA Isolation Method Is a Source of Global DNA Methylation Variability Measured with LUMA. Experimental Analysis and a Systematic Review. PLoS One 8:1–8. 10.1371/journal.pone.0060750

31. Martin M (2013) Cutadapt removes adapter sequences from high-throughput sequencing reads. EMBnet.journal 7:2803–2809. 10.14806/ej.17.1.200

32. Krueger F, Andrews SR (2011) Bismark: a flexible aligner and methylation caller for Bisulfite-Seq applications. Bioinformatics 27:1571–2. 10.1093/bioinformatics/btr167

33. Langmead B, Salzberg SL (2012) Fast gapped-read alignment with Bowtie 2. Nat Methods 9:357–359. 10.1038/nmeth.1923

34. Li H, Handsaker B, Wysoker A, et al (2009) The Sequence Alignment/Map format and SAMtools. Bioinformatics 25:2078–2079. 10.1093/bioinformatics/btp352

35. Ewels P, Magnusson M, Lundin S, Käller M (2016) MultiQC: Summarize analysis results for multiple tools and samples in a single report. Bioinformatics 32:3047–3048. 10.1093/bioinformatics/btw354

36. DI Tommaso P, Chatzou M, Floden EW, et al (2017) Nextflow enables reproducible computational workflows. Nat Biotechnol 35:316–319. 10.1038/nbt.3820

37. New England Biolabs I (2020) NEBNext Enzymatic Methyl-seq Kit. 1–32

38. Chen Y, Pal B, Visvader JE, Smyth GK (2017) Differential methylation analysis of reduced representation bisulfite sequencing experiments using edgeR. F1000Research 6:1–42. 10.12688/f1000research.13196.1

39. Garafutdinov RR, Galimova AA, Sakhabutdinova AR (2019) The influence of CpG (5′-d(CpG)-3′ dinucleotides) methylation on ultrasonic DNA fragmentation. J Biomol Struct Dyn 37:3877–3886. 10.1080/07391102.2018.1533888

40. Hidvégi N, Gulyás A, Dobránszki J (2022) Ultrasound, as a hypomethylating agent, remodels DNA methylation and alters mRNA transcription in winter wheat (Triticum aestivum L.) seedlings. Physiol Plant 1–16. 10.1111/ppl.13777

41. NEB (2023) PERFORMANCE DATA NEBNext UltraShear<sup>TM</sup>

42. Ewels PA, Peltzer A, Fillinger S, et al (2020) The nf-core framework for community-curated bioinformatics pipelines. Nat Biotechnol 38:276–278. 10.1038/s41587-020-0435-1

